# LeafyResNet: Fusarium Wilt Detection in Lettuce Using UAV RGB Imaging and Advanced Deep Learning Model

**DOI:** 10.1101/2025.05.21.655416

**Authors:** Kabir Hossain, Stephanie Slinski, Alexander Bucksch

## Abstract

Lettuce, one of the most consumed leafy greens globally, offers significant health benefits due to its high vitamin, mineral, and fiber content. However, Fusarium wilt, a soil-borne fungus, threatens lettuce yields by reducing both quality and quantity. Traditional disease detection methods, such as manual inspection, are time-consuming and inefficient. This study proposes a Unmanned Aerial Vehicle (UAV)-based approach for detecting Fusarium wilt in lettuce using high-resolution Red-Green-Blue (RGB) imagery. (1) a high resolution RGB lettuce dataset captured by drones at approximately 10 m altitude in collaboration with the Yuma Center of Excellence for Desert Agriculture, (2) identification of candidate Fusarium-infected regions by evaluating 300×300 pixel image patches for light tan coloration, followed by the application of a customized Residual Neural Network (ResNet), called LeafyResNet, to confirm Fusarium presence, and (3) a method for quantifying Fusarium infection severity, which was validated against an expert-ground truth. Our approach to detect Fusarium wilt achieves 96.30% accuracy, 94.10% precision, 100% recall, and a 97.10% F1-score, with a 4% false positive rate. Disease severity scores showed an overall accuracy of 86%. We compared the model to state-of-the-art models, including two variants of ResNet (ResNet18 and ResNet34), Inception_v3, and VGG16. LeafyResNet showed superior results compared to available standard models, highlighting the potential of customizing models for agricultural applications. LeafyResNet provides an efficient and scalable solution for Fusarium wilt monitoring for lettuce crops to advance precision agriculture.

## 1. Introduction

Lettuce (*Lactuca sativa*), a globally consumed leafy vegetable, is valued for its low calorie content and rich nutritional profile, including fiber, iron, folate, and vitamin C (Kim, Moon, Tou, Mou and Waterland, 2016). In the United States, it represented nearly 20% of the $21.8 billion vegetable and melon market in 2022 (US Department of Agriculture, 2020). However, lettuce production is increasingly challenged by factors such as water scarcity, pest pressure, and soil nutrient depletion (Scott, Gordon, Kirkpatrick, Koike, Matheron, Ochoa, Truco and Michelmore, 2012), all of which contribute to crop vulnerability. One significant threat is *Fusarium oxysporum* f. sp. *lactucae*, the causal agent of Fusarium wilt, which severely affects lettuce health and yield.

Traditional disease detection approaches—primarily reliant on manual inspection — are labor-intensive and often reactive, resulting in widespread infection before mitigation can be applied (Sankaran, Mishra, Ehsani and Davis, 2010). Consequently, there is a critical need for scalable, accurate, and early-stage detection methods (Dang, Piran, Han, Min and Moon, 2019).

Recent advancements in remote sensing technologies have facilitated rapid, non-invasive disease monitoring in agriculture (Ferentinos, 2018; Hossain, Villebro and Forchhammer, 2020; Das, Pathan, Jim, Kabir and Mridha, 2025; Bedi and Gole, 2021; V, Bhagwat and Laxmi, 2024). Low-altitude platforms, such as Unmanned Aerial Vehicles (UAVs), offer an affordable and precise means of acquiring high-resolution imagery for crop analysis (Huang, Reddy, Fletcher and Pennington, 2018). Combined with spectral imaging and machine learning, UAVs enable real-time assessment of plant health and early disease detection (Dang et al., 2020a; Leite et al., 2024; Antwi et al., 2024). Detecting Fusarium wilt in lettuce via aerial imagery presents unique challenges. The disease often manifests as subtle yellow-tan discoloration closely resembling the surrounding soil, complicating traditional image-based classification (Wang, Polder, Focker and Liu, 2024). Previous efforts using conditional generative adversarial networks (cGANs) (Isola, Zhu, Zhou and Efros, 2017) for soil segmentation (Li, Stylianou and Pless, 2019) and U-Net architectures (Kugelman, Allman, Read and et al., 2022) for Fusarium wilt detection (Sudars, Namatevs, Nikulins, Balass, Peter, Strautina, Kaufmane and Kalnina, 2023) have struggled with misclassifying infected plants as soil due to color similarity. This problem is particularly relevant in the Sonoran Desert near Yuma, Arizona, which produces approximately 90% of U.S. lettuce from November through April. Here, the soil often shares visual properties with symptomatic leaves—particularly in the light tan color range (RGB: 225, 220, 160). To address this, we trained a deep learning model to emphasize variation within this spectral range, improving the discrimination of infected plant tissue from background soil. In this study, we introduce **LeafyResNet**, a novel deep learning framework designed for accurate detection and severity estimation of Fusarium wilt in lettuce using UAV-acquired RGB imagery. Our method segments imagery into 300×300 pixel patches to localize analysis, enabling the model to capture fine-grained patterns and color variations associated with disease onset. Leveraging the strengths of ResNet’s residual learning architecture (He, Zhang, Ren and Sun, 2016; Rizaldi, Gautama and Kamelia, 2023), LeafyResNet effectively learns complex, nonlinear features from localized image data. Additionally, the model estimates a **Fusarium Infection Severity Index (FISI)**, which quantifies disease severity on a scale from 0 (healthy) to 4 (severely infected), based on expert-labeled data from the Yuma Center of Excellence for Desert Agriculture.

The key contributions of this work include:

- **The first automated method** for detecting Fusarium wilt in lettuce using deep learning and UAV imagery.
- **A novel labeling approach** that assigns severity scores within infected regions, facilitating fine-grained disease monitoring.
- **Light tan intensity analysis** to enhance early detection by distinguishing disease from similar soil tones.
- **A performance comparison** of LeafyResNet against established architectures (ResNet18, ResNet34, VGG16, Inception_v3), revealing improvements in both accuracy and robustness.

The remainder of this paper is organized as follows. Section 2 details the lettuce dataset and preprocessing pipeline. Section 3 describes the patch-based detection framework, including the development of the proposed LeafyResNet model and FISI estimation method. Section 4 presents experimental results, model evaluations, and a comparison with existing approaches. Section 5 concludes the study and outlines directions for future research.

## 2. Lettuce Dataset Description

### 2.1. Data Acquisition

Data for this study were collected using a DJI Inspire 1 drone equipped with an RGB camera in a 2-acre field in Yuma County, Arizona. Flights were carried out by trained staff at the Yuma Center of Excellence for Desert Agriculture. A total of 6,239 images were obtained from two adjacent fields: the north field (2,704 images) and south field (3,535 images). The data collection spanned nine weeks, covering two regions within the field, with a total of 6,239 images captured from lettuce plants of various genotypes. Data acquisition began approximately one month after planting, with weekly drone flights from October 11 to December 13, 2023. No data was collected during the seventh week due to adverse weather. The drone captured high-resolution images from an altitude of approximately 10 meters with a resolution of 4000 × 3000 pixels. The altitude of 10 meters was chosen to obtain images with a clear and broad view of the lettuce field. At this altitude, the images have sufficient resolution for detecting wilting while still covering a larger portion of the field. Details of the RGB lettuce dataset are summarized in Table 1. Fusarium wilt, initially absent, began spreading in early November. By November 2, 10.23% of images were affected, with a sharp increase in the north field to 19.76% by November 7. Manual inspection found that approximately 44.85% of the images in both fields showed at least one wilting plant after nine weeks. An example of a UAV (drone) image of the lettuce field with highlighted Fusarium wilt symptoms is shown in Figure 1.

**Table 1.** Details of the RGB lettuce dataset collected in Yuma County, Yuma, Arizona, using the DJI Inspire 1 drone at an altitude of 10 meters. The dataset includes images from two adjacent fields: the North field with 2,704 images and the South field with 3,535 images, totaling 6,239 images.

| Region (Yuma County, Arizona) | Date | #Fields | Altitude (m) | #RGB Images |
| --- | --- | --- | --- | --- |
| North field | Oct 11 - Dec 13, 2023 | 1 | 10 | 2,704 |
| South field | Oct 11 - Dec 13, 2023 | 1 | 10 | 3,535 |
| Total |  | 2 | 10 | 6,239 |

**Figure 1:**
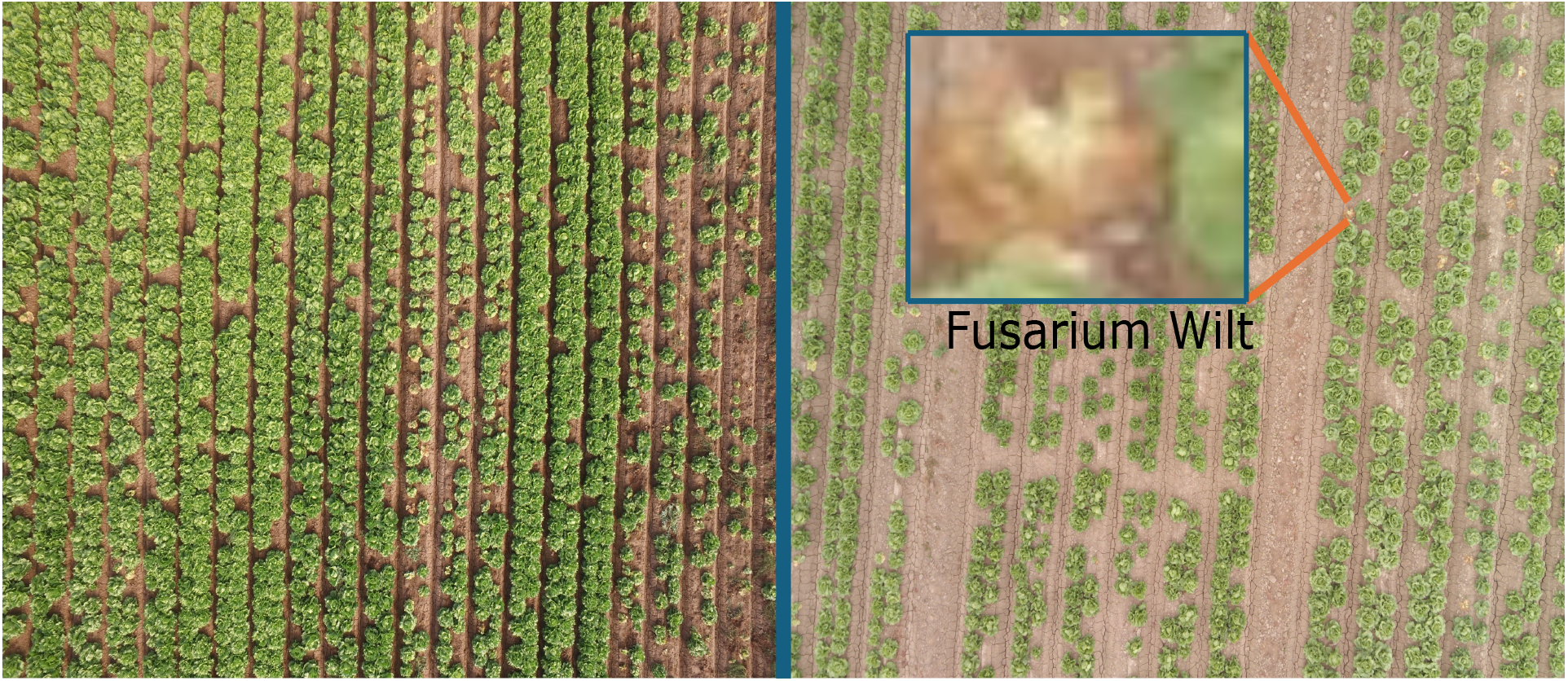
UAV-captured RGB image and highlighted Fusarium wilt symptoms in lettuce field. Left: A high-resolution RGB image captured by a DJI Inspire 1 drone at an altitude of approximately 10 meters, showing the overall lettuce field. Right: A full image of the lettuce with Fusarium wilt symptoms, where areas affected by the disease are highlighted for clarity. The highlighted areas emphasize the typical yellowing and wilting of lettuce leaves due to Fusarium wilt, as observed during the study.

### 2.2. Training Dataset Preparation

For training the LeafyResNet model, the dataset first needed to be prepared. A total of 30 full RGB images were selected from the north field, from which 675 image patches (100 × 100 pixels) were extracted, representing both healthy and Fusarium-infected regions. These Region of Interest (ROI) images were preprocessed to ensure consistency and improve model performance. Specifically, pixel values were normalized by scaling the pixel intensities to the range [0, 1], which helps the model converge faster during training.

The ROI patches (see Figure 2) were then split into an 80:20 ratio for training and testing, with the training set containing 540 ROI images and the testing set containing 135 images. Data augmentation was applied only to the training set, expanding it to 57,200 image patches, while the testing set remained non-augmented to ensure a fair evaluation of the model on original, unseen data.

**Figure 2:**
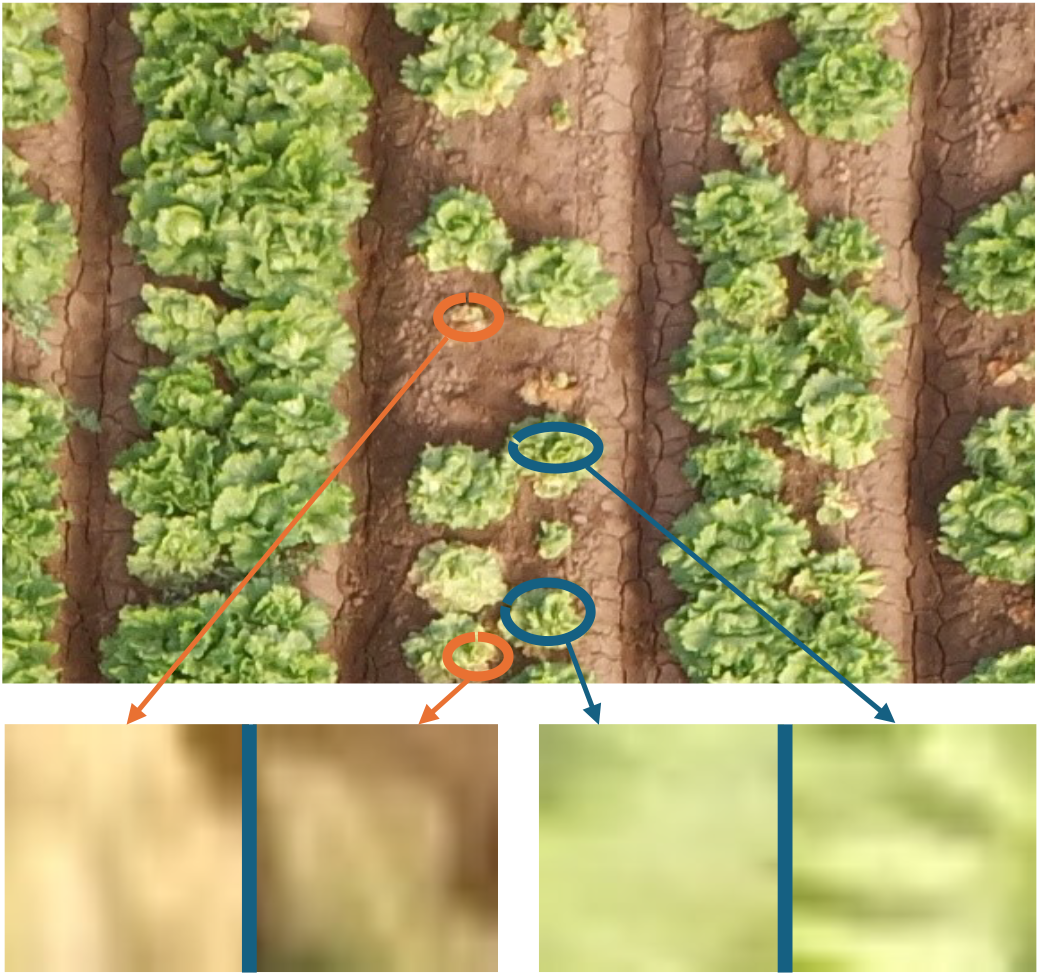
An RGB image of a lettuce sample captured by UAV, showing two distinct ROI types: blue circles indicate healthy lettuce regions, while orange circles mark Fusarium-infected ROIs.

**Figure 3:**
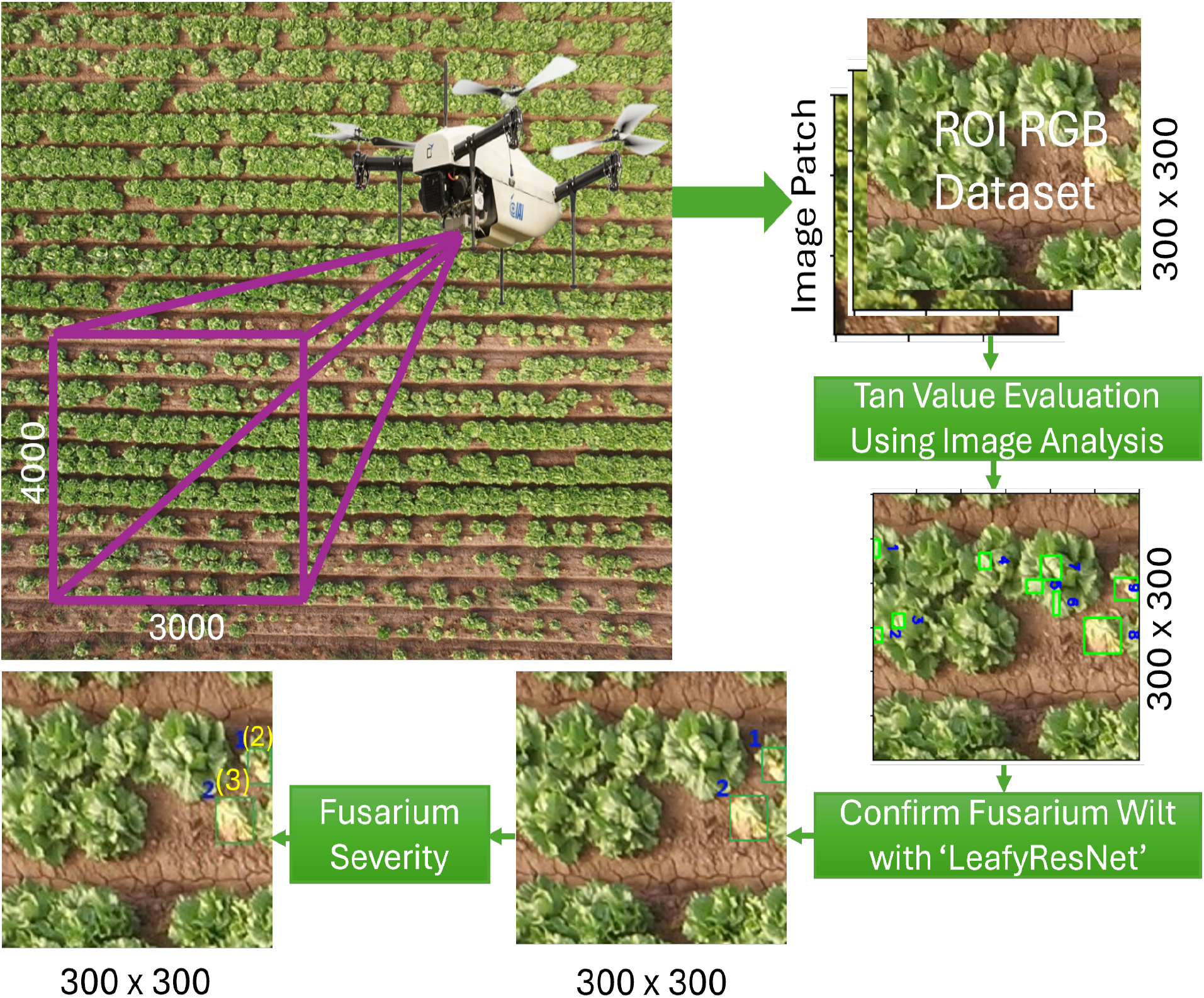
Proposed methodology for Fusarium wilt detection in lettuce using UAV-based RGB imagery. Steps include Region of Interest (ROI) RGB dataset preparation from the full image, light tan intensity evaluation, classification with the LeafyResNet model, and severity quantification by labeling for effective agricultural monitoring.

The data augmentation techniques used in this study included rotation, translation, shearing, zooming, and horizontal flipping. Rotation and flipping (Huang, Pan, Zhang, Qian, Gao and Wu, 2019) were applied to introduce variability in image orientation. Shearing and zooming (Grochowski et al., 2018) were used to simulate distortions and scale variations, while translation (Engstrom, Tsipras, Schmidt and Madry, 2017) was implemented to shift images spatially, creating a more diverse dataset and improving model generalization.

After augmentation, the LeafyResNet model was trained to classify two categories: healthy and Fusarium wilt, confirming the presence of Fusarium or healthy tissue in each region.

## 3. Methodology

### 3.1. Patch Creation

The patch-based image method to detect Fusarium wilt in lettuce is inspired by techniques used in agricultural image analysis. Previous work by Hasan et al. (2023) on crop and weed recognition through patch-based deep learning, and Moazzam et al. (2021) on weed detection in sugar beet using patch-image classification, demonstrate improved accuracy using patches for localized feature analysis. This segmentation improves the model’s ability to evaluate finer details within an image, which is a crucial aspect in plant disease detection because small, localized changes in texture or color are important indicators of disease.

To implement this localized analysis for the detection of Fusarium wilt, our method divides the original image, which has a resolution of 4000 × 3000 pixels (4K resolution), into smaller patches of 300 × 300 pixels.

For an original image with dimensions *H* × *W* (height × width) and a patch size of *S*_*x*_ × *S*_*y*_ (300 × 300 pixels), the total number of patches *T*_*x*_ and *T*_*y*_ along he width and height are calculated as:

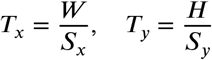

The image will then be divided into a grid of 13 × 10 patches, as the original 4000 × 3000 image is divided by the 300 × 300 patch size (4000/300 = 13 and 3000/300 = 10). Each patch *p*_*i,j*_ is extracted from the image starting at coordinates (*x*_0_, *y*_0_), where:

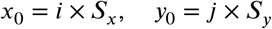

*S*_*x*_ and *S*_*y*_ are set to 300, making each patch 300 × 300 pixels. These patches are used to estimate the light-tan color range, which aids in the identification of the candidate Fusarium infection.

### 3.2. Fusarium Wilt Detection Using Deep Learninig

#### 3.2.1. Detection of Fusarium infected candidate regions

We evaluate the light tan color value as a feature for candidate Fusarium wilt detection. The process begins by evaluating the image patches created earlier for further processing. We sampled 50 image patches for soil color samples and generated a histogram. The median values of the histogram were calculated as R = 225, G = 208, and B = 160, which fall within the tan color range. Since Fusarium appears lighter than these values, we selected candidate pixels of Fusarium wilt as equal to or greater than the median values. After that, a mask is generated to isolate candidate regions of connected pixels. To further refine the mask and improve the detection of contours (OpenCV team), image processing techniques are applied. Morphological operations, including dilation and erosion, are used to improve the precision of contour identification. Dilation expands the detected regions, merging neighboring areas that may have been separated because of noise or inconsistencies in the image. Erosion is then applied to reduce the size of the expanded regions, eliminating small, irrelevant features, and sharpening the boundaries of the contours.

Finally, the contours are extracted from the processed image, to identify candidate areas of Fusarium wilt. This approach leverages the distinctive yellow-brown discoloration of infected plants, as well as the potential to efficiently identify Fusarium from soil color, which may resemble the color of the disease. Using contour detection, a rectangle box is created around each candidate region.

#### 3.2.2. Implementation of the LeafyResNet Model

We used LeafyResNet to confirm candidate regions of Fusarium wilt. The model begins with an initial convolutional layer using a 7×7 kernel and 64 filters, followed by batch normalization and a ReLU activation function.

A max-pooling layer with a 3×3 kernel and stride 2 is then applied to reduce the spatial dimensions. The core of the model consists of several residual blocks. Each block includes two convolutional layers with a 3×3 kernel, batch normalization, and ReLU activations. The residual connections within each block add the input to the output of the convolutions, helping the model learn residual mappings and mitigating the problem of vanishing gradients (Hochreiter, 1998), thereby improving training efficiency. The architecture of the LeafyResNet model is detailed in Table 2.

**Table 2.** Layer-wise Architecture of ResNet for Fusarium Confirmation in Lettuce Crops.

| Layer Type | Output Shape (Size) | Description |
| --- | --- | --- |
| Input Layer | (100, 100, 3) | Input image with 3 color channels (RGB) |
| Conv1 | (100, 100, 64) | 7x7 convolution, 64 filters, stride=2, BatchNorm, ReLU |
| MaxPooling | (50, 50, 64) | MaxPooling (3x3), stride=2 to reduce spatial dimensions |
| Residual Block 1 | (50, 50, 64) | 2x3x3 convolution, BatchNorm, ReLU, skip connection |
| Residual Block 2 | (50, 50, 64) | 2x3x3 convolution, BatchNorm, ReLU, skip connection |
| Residual Block 3 | (50, 50, 64) | 2x3x3 convolution, BatchNorm, ReLU, skip connection |
| Residual Block 4 | (25, 25, 128) | 2x3x3 convolution, BatchNorm, ReLU, stride=2, skip connection |
| Residual Block 5 | (25, 25, 128) | 2x3x3 convolution, BatchNorm, ReLU, skip connection |
| Residual Block 6 | (25, 25, 128) | 2x3x3 convolution, BatchNorm, ReLU, skip connection |
| Residual Block 7 | (13, 13, 256) | 2x3x3 convolution, BatchNorm, ReLU, stride=2, skip connection |
| Residual Block 8 | (13, 13, 256) | 2x3x3 convolution, BatchNorm, ReLU, skip connection |
| Residual Block 9 | (13, 13, 256) | 2x3x3 convolution, BatchNorm, ReLU, skip connection |
| Residual Block 10 | (7, 7, 512) | 2x3x3 convolution, BatchNorm, ReLU, stride=2, skip connection |
| Residual Block 11 | (7, 7, 512) | 2x3x3 convolution, BatchNorm, ReLU, skip connection |
| Residual Block 12 | (7, 7, 512) | 2x3x3 convolution, BatchNorm, ReLU, skip connection |
| Global Average Pooling | (1, 1, 512) | Reduces the spatial dimensions to 1x1 |
| Dense (Output) | (1,) | Fully connected layer, sigmoid activation for binary classification |

### 3.3. Estimation of Fusarium Infection Severity Index (FISI)

Following the confirmation of Fusarium wilt regions by the LeafyResNet model, we hypothesize that the brightness of the pixels in the identified regions can be used to estimate FISI in the detected area. Specifically, we assume that regions with brighter pixels correspond to a more severe infection. To estimate infection severity, we perform a pixel-level analysis by converting the identified regions from RGB to the HSV color space, where *H* represents the Hue, *S* represents the Saturation, and *V* represents the Value, which is the brightness component.

The use of HSV for this analysis is motivated by its ability to separate image intensity from color information, making it more robust to lighting variations and shadow effects compared to the RGB color space. This approach has already been validated (Dang et al., 2020a). Here, the HSV color space was effective to analyze the severity of plant disease, particularly in outdoor conditions where consistent illumination cannot be guaranteed (Huang et al., 2018). In our case, this method helps to isolate brightness variations that are indicative of Fusarium infection, which might otherwise be obscured in the RGB space due to color shifts.

We used the cv2.cvtColor function from OpenCV (OpenCV Team) to convert the ROI from RGB to the HSV color space. Once the images are converted to the HSV color space, we apply a threshold to the Value channel (*V*) to isolate bright pixels, which are presumed to represent the Fusarium-infected areas. The thresholding operation is defined as:

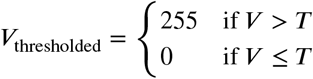

where *V* is the pixel intensity in the Value channel, and *T* is the threshold value, estimated by analyzing randomly selected Fusarium-infected regions across 30 full images with a resolution of 4000 × 3000. To estimate *T*, a histogram is generated from these selected regions, and the mean value of the histogram is calculated. This mean value is used as the threshold *T*.

Mathematically, the threshold *T* is computed as the average of the pixel intensities in the Value channel from the usarium-infected regions:

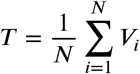

where *N* is the total number of pixels belonging to detected Fusarium regions, and *V*_*i*_ represents intensities across the individual pixels i in the Value channel.

After thresholding, we count the number of bright pixels (*N*_bright_) and calculate the total number of pixels in the egion (*N*_total_). The severity of the disease is quantified as the percentage of bright pixels:

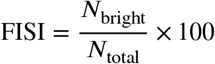

Where:

- *N*_bright_ is the number of bright pixels (those with *V* > *T*),
- *N*_total_ is the total number of pixels in the ROI.

After quantifying Fusarium wilt severity as a percentage, labels are assigned based on the following thresholds: 0 ≤ value < 40 for Category 1, 40 ≤ value < 55 for Category 2, 55 ≤ value < 65 for Category 3, and 65 ≤ value ≤ 100 for Category 4.

## 4. Experimental Results

### 4.1. Performance Analysis of Fusarium Wilt Detection

#### 4.1.1. Candidate Fusarium Wilt Detection Results

We identified candidate Fusarium-infected regions by detecting the specific light tan color range. Overall, we visually identified 68 candidate Fusarium-infected areas across 10 image patches. Of these 68 candidate areas, 63 were correctly identified by our model, while 5 were misclassfied. Additionally, 40 non-Fusarium regions were incorrectly marked as candidate Fusarium-infected, reflecting the model’s sensitivity but also indicating the need for further validation. Despite these challenges, the model showed strong performance with an overall precision of 92.65%. In severe Fusarium cases, where the infection color was very close to the soil color, the model struggled to distinguish them, leading to missed detections. However, excluding these severe cases, the precision significantly improved, surpassing 99%. This highlights the model’s ability to accurately identify infections in early-stage Fusarium infections.

In Figure 5 (a-b), we visualize the intermediate candidate Fusarium wilt detection results, where the identified regions are highlighted by rectangles. These “candidate regions” represent areas that may show Fusarium-like discoloration, but do not confirm the presence of Fusarium wilt. Further confirmation is required, which is addressed in the next section using the proposed LeafyResNet model.

#### 4.1.2. Fusarium Wilt Confirmation and Performance

The proposed LeafyResNet model demonstrated strong performance in confirming Fusarium infections within the candidate areas. The training performance of the proposed model is shown in Figure 4, where (a) illustrates the loss curve over epochs, and (b) displays the accuracy progression during training. These plots provide insights into how the model converges over time, highlighting its efficiency in both, minimizing loss and maximizing accuracy. Evaluation metrics on the test set for the proposed LeafyResNet model are summarized in Table 3, showing an overall accuracy of 96.3%, precision of 94.1%, recall of 100%, F1-score of 97.1%, along with a False Positive Rate (FPR) of 4%. Figure 5 (b) illustrates the candidate Fusarium-infected regions detected using light tan value evaluation, highlighted by rectangles. These regions were identified based on light tan discoloration intensity but included false positives due to visual similarities with infected areas, which caused confusion in the detection process. Figure 5 (c) shows the confirmed Fusarium-infected areas after applying the LeafyResNet model. The model effectively removed these false positives, retaining only the true infections. This two-step process improved accuracy by confirming the initially detected Fusarium infections.

**Table 3.**
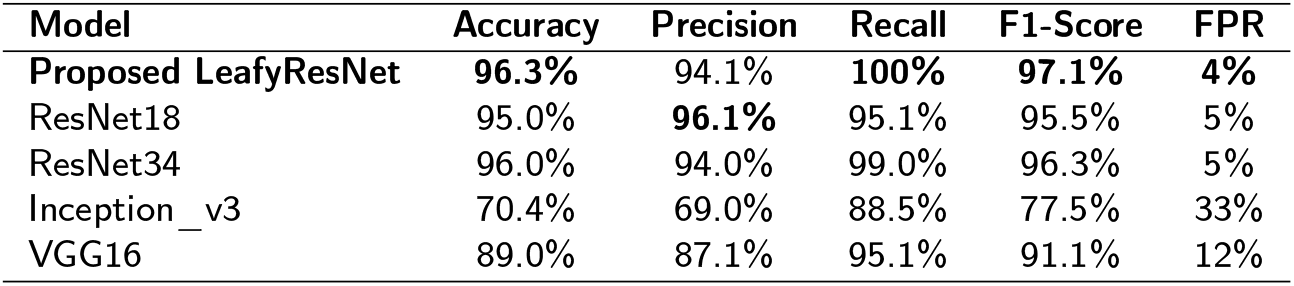
Performance comparison of different models on the lettuce dataset.

| Model | Accuracy | Precision | Recall | F1-Score | FPR |
| --- | --- | --- | --- | --- | --- |
| **Proposed LeafyResNet** | **96.3%** | 94.1% | **100%** | **97.1%** | **4%** |
| ResNet18 | 95.0% | **96.1%** | 95.1% | 95.5% | 5% |
| ResNet34 | 96.0% | 94.0% | 99.0% | 96.3% | 5% |
| Inception_v3 | 70.4% | 69.0% | 88.5% | 77.5% | 33% |
| VGG16 | 89.0% | 87.1% | 95.1% | 91.1% | 12% |

**Figure 4:**
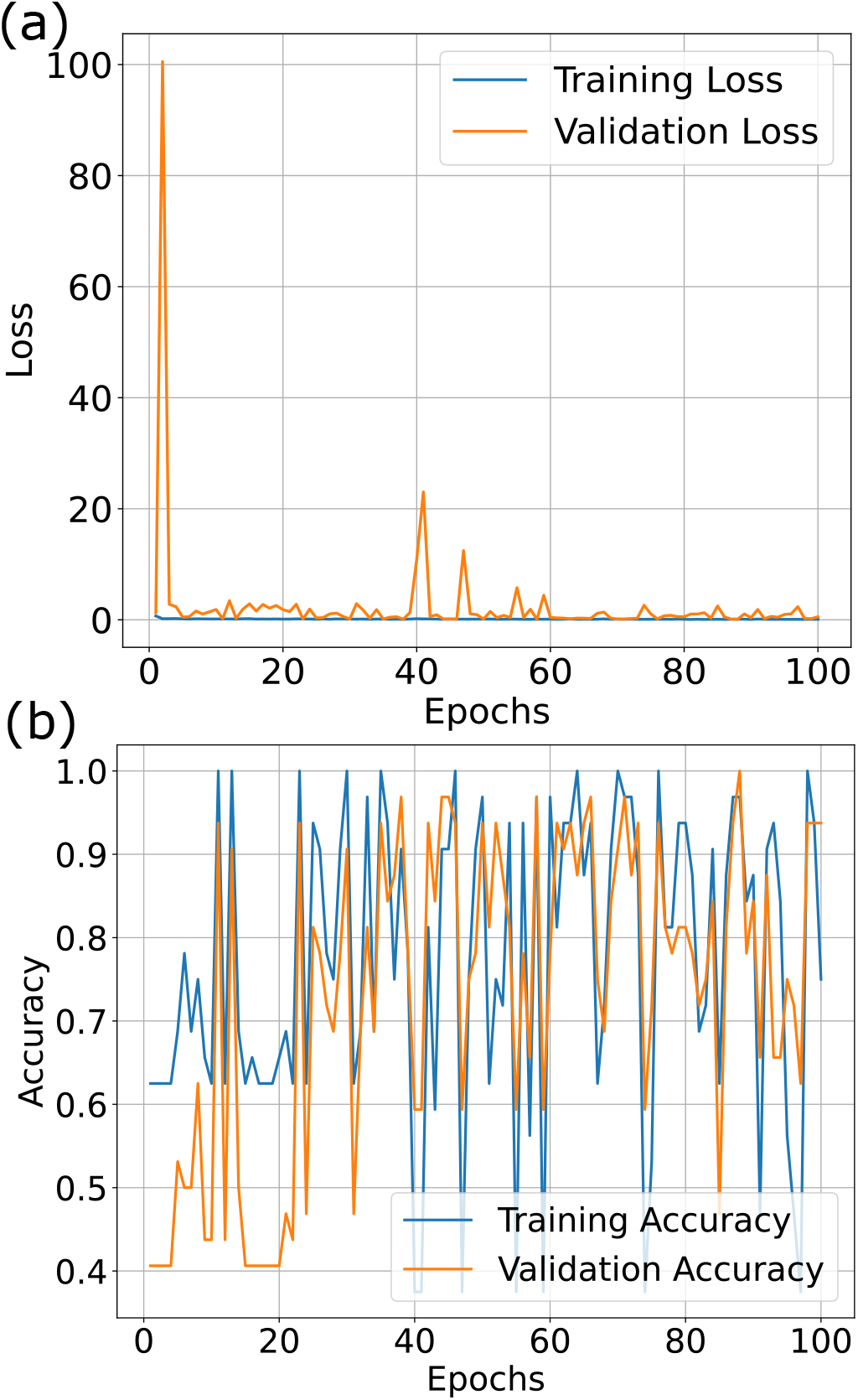
Training Performance: (a) Loss vs Epochs and (b) Accuracy vs Epochs for the Proposed Customized ResNet Model.

**Figure 5:**
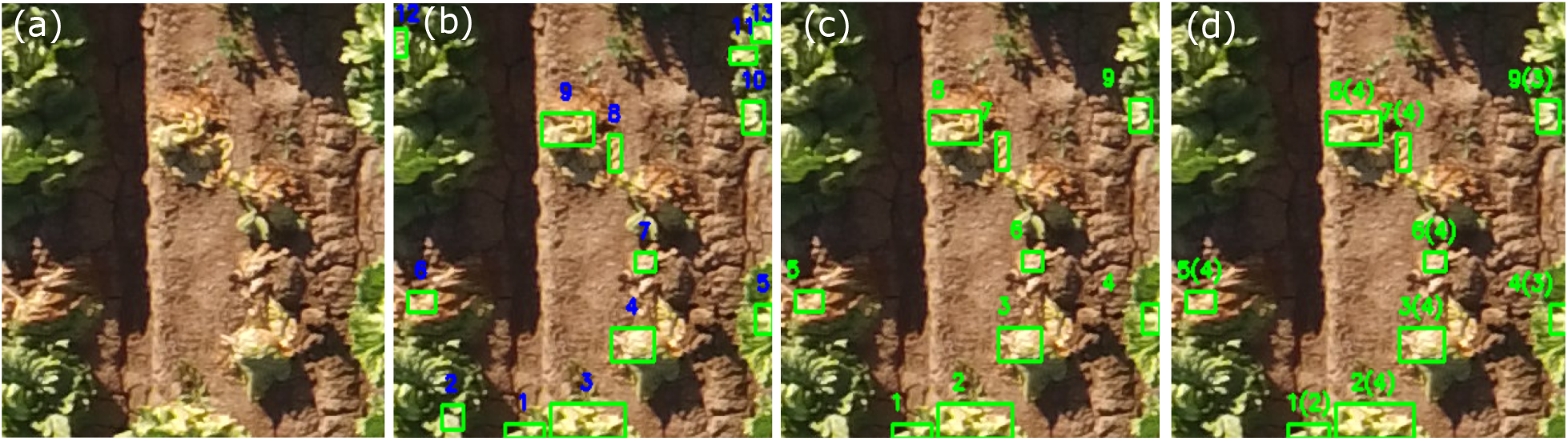
Workflow of Fusarium Wilt Detection and Severity Quantification in Lettuce. (a) Original Image (b) candidate Fusarium Wilt Detection (c) Confirm Fusarium Wilt, and (d) Quantification of Severity in detected regions, with each detected region labeled by a number followed by its severity in parentheses, e.g., 1 (4) indicating detected region 1 with severity level 4.

#### 4.1.3. Performance Comparison with State-of-the-Art Models

To evaluate the effectiveness of the proposed model, we compared its performance with state-of-the-art deep learning models using the proposed lettuce dataset. The first benchmark model, Inception-V3 (Dang, Ibrahim Hassan, Suhyeon, kumar Sangaiah, Mehmood, Rho, Seo and Moon, 2020b), was originally designed to distinguish between healthy and Fusarium-infected regions in radish. The second model, VGG-16 (Ha, Moon, Kwak, Hassan, Dang, Lee and Park, 2017), was previously used to classify healthy and wilted radish plants. We implemented these models on our lettuce dataset for a direct comparison. Additionally, we included two ResNet variants (ResNet18 (He et al., 2016) and ResNet34 (Rizaldi et al., 2023)), to further analyze model performance.

A comprehensive performance comparison was conducted between the proposed LeafyResNet model, Inception-V3,VGG-16, and the two ResNet variants (ResNet18 and ResNet34). All models were trained on the same lettuce dataset to ensure a fair evaluation. The results are summarized in Table 3. The best results for each metric are highlighted in bold for clarity.

The proposed LeafyResNet model outperforms the comparison models in terms of overall performance, achieving 96.3% accuracy, 94.1% precision, 100% recall, 97.1% F1-score, and a low False Positive Rate (FPR) of 4%. Although ResNet18 achieved a slightly higher precision of 96.1%, LeafyResNet excels in accuracy, recall, and F1-score. Additionally, it significantly reduces false positives compared to other models, ensuring more reliable and robust results. While ResNet34, Inception_v3, and VGG16 show good performance, they fall short in comparison to the performance and efficiency of the LeafyResNet model, demonstrating its potential for real-world applications.

### 4.2. Performance Evaluation of FISI

The model was further tested for its ability to quantify the severity of Fusarium infections in lettuce crops. By analyzing the pixel composition within the confirmed infected regions, we assessed the severity of the infection. Fusarium wilt severity was quantified by assigning labels (0 for healthy, 1–4 for varying severity) in the detected Fusarium wilt regions, assigning non-detected areas as 0. The quantification process evaluated the distribution of labels within the identified regions, with higher label values representing more severe infections. For example, the confirmed regions in Figure 5 (d) show nine detected areas, each enclosed in a bounding box and labeled with its box number followed by its severity level in parentheses (e.g., 1(2), 2(4)), demonstrating the model’s ability to distinguish different infection severity levels. To evaluate the model we used severity data collected manually in the the south field. Ground truth labels (0–4) were assigned to each plant by experts at the Yuma Center of Excellence for Desert Agriculture, and 10 randomly selected patches containing 112 plants were used for evaluation. The model achieved an overall accuracy of 86%, performing best for Class 0 (96.3% accuracy, 50 plants). For infected plants, accuracy was 80% for Class 1 (10 plants), while Classes 2–4 ranged from 69% to 76% accuracy across 67 plants. The model demonstrated reliability to identify FISI, despite some overlap in the classification of severity stages, particularly between Class 3 and Class 4, and Class 2 and Class 3.

### 4.3. Discussion

The proposed method for detecting Fusarium wilt and scoring Fusarium infection severity demonstrated improved performance compared to state-of-the-art methods but also encountered challenges. By integrating RGB thresholding for initial detection with the deep learning model LeafyResNet, we achieved high accuracy (96.3%), precision (94.1%), recall (100%), and an F1-score of 97.1%, with a false positive rate of 4%. However, detection accuracy declined particularly at advanced infection stages, when the color of Fusarium-infected areas became similar to that of the soil.

However, emphasizing the light tan color during model training alleviated some of the previously reported limitations (e.g., (Dang et al., 2020a)), although some detection challenges remain when color differences become minimal. These challenges arise when severely infected Fusarium lightens, causing pixel values to decrease and become too similar to the soil average color range (216, 200, 153).

It is also important to note that Fusarium wilt can resemble other wilting conditions, such as those caused by water stress or root damage due to insect feeding. While our evaluations were conducted in a field where Fusarium wilt was the only disease present, the presence of other pests or diseases in the field could lead to misidentifying other diseases as Fusarium wilt. Incorporating these additional diseases into the training data could help distinguish Fusarium wilt from other similar conditions.

Notably, our model excelled in early stage detection, achieving a precision of 92.65%, which increased to over 99% when severe cases were excluded. Comparative analysis with state-of-the-art models, including Inception_v3, VGG16, ResNet18, and ResNet34, confirmed the superior performance of our proposed LeafyResNet model in detecting Fusarium wilt in different datasets. Future improvements could integrate environmental metadata to enhance model robustness.

The severity quantification of our model achieved an overall accuracy of 86% against expert-labeled ground truth data from the Yuma Center of Excellence for Desert Agriculture. However, challenges in plant-to-plant matching due to time differences between UAV image collection and visual disease scoring affected classification accuracy. Careful adjustments during validation helped mitigate these discrepancies, ensuring reliable severity assessments. Expanding this approach to diverse crops and integrating real-time drone-based detection could further enhance its applicability in precision agriculture, improving disease monitoring and management strategies.

From a scientific standpoint, our method also contributes to advancements in root biology. Fusarium infection originates at the root, and root architecture is one potential indicator of a plant’s tolerance to biotic stressors, such as the parasitic weed Striga, particularly in crops like cowpea (Burridge, Schneider, Huynh, Roberts, Bucksch and Lynch, 2017) and sorghum (Kawa, Thiombiano, Shimels, Taylor, Walmsley, Vahldick, Rybka, Leite, Musa, Bucksch et al., 2024; Kawa, Taylor, Thiombiano, Musa, Vahldick, Walmsley, Bucksch, Bouwmeester and Brady, 2021). While a direct pathogenic comparison between Striga and the soil-borne fungus Fusarium is not feasible due to their differing infection mechanisms, both interact directly with the root system. This suggests that phenotyping root architectural traits could play a pivotal role in understanding and potentially enhancing plant tolerance to Fusarium infection using modern 3D root phenotyping tools ((Liu, Barrow, Hanlon, Lynch and Bucksch, 2021; Liu, Bonelli, Pietrzyk and Bucksch, 2023)) by associating Fusarium severity and root architecture phenotype.

## 5. Conclusion

We have developed a highly effective method for detecting Fusarium wilt in lettuce and accurately estimating the FISI using UAV imagery. By integrating light tan evaluation for initial detection with a deep learning model featuring the LeafyResNet architecture, our approach delivers exceptional performance, achieving test results of 96.3% accuracy, 94.1% precision, 100% recall, and a 97.1% F1-score, with a false positive rate of just 4%, outperforming the state-of-the-art models, including Inception_v3, VGG16, ResNet18, and ResNet34. Moreover, our model demonstrated an 86% overall accuracy in assessing Fusarium infection severity, which was validated against expert-annotated ground truth.

Critically, our approach excels at identifying early-stage symptoms of Fusarium wilt in lettuce, enabling timely intervention to mitigate crop losses. By detecting infections at different stages of progression, our model not only enhances disease monitoring but also provides precise quantification of infection severity across affected regions. These capabilities make it a promising tool for precision agriculture and can be integrated into state-of-the-art data processing pipelines used for lettuce (Gonzalez, Zarei, Hendler, Simmons, Zarei, Demieville, Strand, Rozzi, Calleja, Ellingson et al., 2023). Therefore, the presented method offers growers reliable, data-driven insights to optimize disease management strategies.

Looking ahead, expanding this methodology to other crops and plant diseases will enhance its applicability across diverse agricultural landscapes. Further advancements in real-time, drone-based detection will empower farmers with instant, actionable intelligence to reduce yield losses, minimize chemical inputs, and promote sustainable farming practices. This aligns with global efforts to combat climate change, enhance food security, and support climate-resilient agriculture.

Additionally, refining our quantification approach to assess infection severity across entire fields—rather than isolated detected regions—will provide a more comprehensive evaluation of disease impact. Such advancements will enable more accurate yield loss estimations and economic impact assessments, further strengthening decision-making in precision agriculture.

In conclusion, our model’s performance underscores its transformative potential for real-world agricultural applications. Future research should refine its ability to distinguish between adjacent severity levels, further enhancing its precision and effectiveness. With its proven capability in early detection and severity assessment, this approach represents a significant step forward in disease monitoring and crop protection.

## Acknowledgment

The work is supported by the start-up package of Dr. Alexander Bucksch from the University of Arizona College of Agriculture, Life and Environmental Sciences. We would also like to acknowledge Rosa Bevington for her invaluable assistance as the drone pilot and for collecting all the data used in this research.

## CRediT authorship contribution statement

**Kabir Hossain:** Writing – original draft, Visualization, Software, Methodology, Investigation, Formal analysis. **Stephanie Slinski:** Writing – review & editing, Supervision, Resources, Data collection, Supervision of Lettuce Cultivation. **Alexander Bucksch:** Conceptualization, Supervision, Funding acquisition, Writing – review & editing.

---

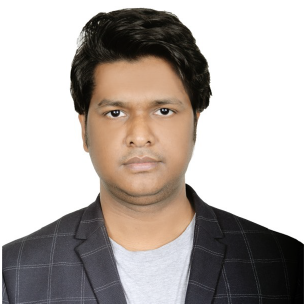

I am a postdoctoral researcher at the School of Plant Sciences, University of Arizona, with experience in machine learning for plant phenotyping, disease detection, underground plant root growth monitoring using AI, precision agriculture using drone imagery, drone image processing, signal processing, data science and computer vision. My research integrates AI and UAV-based remote sensing to advance sustainable farming. I hold a Ph.D. in Electrical and Photonics Engineering from the Technical University of Denmark, where I focused on drone image processing for district heating systems. I earned my B.Eng. in Computer Engineering and M.Eng. in Electronics and Computer Engineering from Chonnam National University, South Korea.

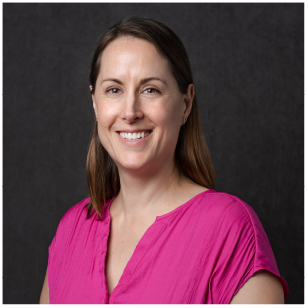

Stephanie Slinski is the Associate Director of Applied Research and Development at the Yuma Center of Excellence for Desert Agriculture (YCEDA), a public-private partnership between the desert agriculture industry and the University of Arizona. In her role she develops research projects, collaborations, partnerships, outreach and education opportunities to support the needs of the desert ag industry including but not limited to, plant health, soil health, Agtech, water conservation, and agriculture sustainability. Before coming to YCEDA in 2018, Stephanie worked with the citrus industry in Florida, mainly focusing on projects addressing citrus greening. She has degrees in Plant and Soil Science, Microbiology and a PhD in Plant Pathology from the University of California, Davis.

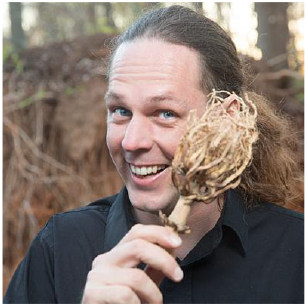

Alexander Bucksch is an Associate Professor in the School of Plant Sciences at the University of Arizona who develops plant phenotyping methods across all biological and ecological scales with an emphasis on plant roots. As a trained computer scientist, he developed his interest in plant biology & ecology during his undergraduate studies at the Brandenburg Technical University. Since then, he developed computational methods to analyze plant morphology in the field as a PhD at the Delft Technical University and as a PostDoc at the Georgia Institute of Technology. He was awarded the NSF CAREER Award, the Fred C. Davison Early Career Award, and the Early Career Award of the North American Plant Phenotyping Network for his computational approaches to understanding the function of natural variation in plant phenotypes.

## References

Antwi, K., et al., 2024. On the application of image augmentation for plant disease detection: A systematic literature review. Smart Agricultural Technology 9, 100590. doi:10.1016/j.atech.2024.100590.

Bedi, P., Gole, P., 2021. Plant disease detection using hybrid model based on convolutional autoencoder and convolutional neural network. Artificial Intelligence in Agriculture 5, 90–101. doi:10.1016/j.aiia.2021.05.002.

Burridge, J.D., Schneider, H.M., Huynh, B.L., Roberts, P.A., Bucksch, A., Lynch, J.P., 2017. Genome-wide association mapping and agronomic impact of cowpea root architecture. Theoretical and applied genetics 130, 419–431.

Dang, L., Piran, J., Han, D., Min, K., Moon, H., 2019. A survey on internet of things and cloud computing for healthcare. Electronics 8, 768. doi:10.3390/electronics8070768.

Dang, L., et al., 2020a. Fusarium wilt of radish detection using rgb and near infrared images from unmanned aerial vehicles. Remote Sens. 12, 2863. doi:10.3390/rs12172863.

Dang, L.M., Ibrahim Hassan, S., Suhyeon, I., kumar Sangaiah, A., Mehmood, I., Rho, S., Seo, S., Moon, H., 2020b. Uav based wilt detection system via convolutional neural networks. Sustainable Computing: Informatics and Systems 28, 100250. URL: https://www.sciencedirect.com/science/article/pii/S2210537917304018, xdoi:10.1016/j.suscom.2018.05.010.

Das, A., Pathan, F., Jim, J.R., Kabir, M.M., Mridha, M., 2025. Deep learning-based classification, detection, and segmentation of tomato leaf diseases: A state-of-the-art review. Artificial Intelligence in Agriculture 15, 192–220. doi:10.1016/j.aiia.2025.02.006.

Engstrom, L., Tsipras, D., Schmidt, L., Madry, A., 2017. A rotation and a translation suffice: Fooling cnns with simple transformations. ArXiv Preprint arXiv:1712.02779. URL: https://arxiv.org/abs/1712.02779.

Ferentinos, K.P., 2018. Deep learning models for plant disease detection and diagnosis. Computers and Electronics in Agriculture 145, 311–318. URL: https://www.sciencedirect.com/science/article/pii/S0168169917311742, xdoi:10.1016/j.compag.2018.01.009.

Gonzalez, E.M., Zarei, A., Hendler, N., Simmons, T., Zarei, A., Demieville, J., Strand, R., Rozzi, B., Calleja, S., Ellingson, H., et al., 2023. Phytooracle: Scalable, modular phenomics data processing pipelines. Frontiers in Plant Science 14, 1112973.

Grochowski, M., et al., 2018. Data augmentation for improving deep learning in image classification problem, in: 2018 International Interdisciplinary PhD Workshop (IIPhDW), pp. 1–5. doi:10.1109/IIPHDW.2018.8388338.

Ha, J.G., Moon, H., Kwak, J.T., Hassan, S.I., Dang, M., Lee, O.N., Park, H.Y., 2017. Deep convolutional neural network for classifying fusarium wilt of radish from unmanned aerial vehicles. Journal of Applied Remote Sensing 11, 042621. doi:10.1117/1.JRS.11.042621.

Hasan, A.M., et al., 2023. Image patch-based deep learning approach for crop and weed recognition. Ecological Informatics 78, 102361. doi:10.1016/j.ecoinf.2023.102361.

He, K., Zhang, X., Ren, S., Sun, J., 2016. Deep residual learning for image recognition, in: Proceedings of the 2016 IEEE Conference on Computer Vision and Pattern Recognition (CVPR), pp. 770–778. doi:10.1109/CVPR.2016.90.

Hochreiter, S., 1998. The vanishing gradient problem during learning recurrent neural nets and problem solutions. International Journal of Uncertainty, Fuzziness and Knowledge-Based Systems 06, 107–116. doi:10.1142/S0218488598000094.

Hossain, K., Villebro, F., Forchhammer, S., 2020. Uav image analysis for leakage detection in district heating systems using machine learning. Pattern Recognition Letters 140, 158–164. URL: https://www.sciencedirect.com/science/article/pii/S0167865520302038, xdoi:10.1016/j.patrec.2020.05.024.

Huang, L., Pan, W., Zhang, Y., Qian, L., Gao, N., Wu, Y., 2019. Data augmentation for deep learning-based radio modulation classification. IEEE Access 8, 1498–1506. doi:10.1109/ACCESS.2019.2960775.

Huang, Y., Reddy, K., Fletcher, R., Pennington, D., 2018. Uav low-altitude remote sensing for precision weed management. Weed Technol. 32, 2–6. Available: 10.1017/wet.2017.89.

Isola, P., Zhu, J.Y., Zhou, T., Efros, A.A., 2017. Image-to-image translation with conditional adversarial networks, in: 2017 IEEE Conference on Computer Vision and Pattern Recognition (CVPR), pp. 5967–5976. doi:10.1109/CVPR.2017.632.

Kawa, D., Taylor, T., Thiombiano, B., Musa, Z., Vahldick, H.E., Walmsley, A., Bucksch, A., Bouwmeester, H., Brady, S.M., 2021. Characterization of growth and development of sorghum genotypes with differential susceptibility to striga hermonthica. Journal of Experimental Botany 72, 7970–7983.

Kawa, D., Thiombiano, B., Shimels, M.Z., Taylor, T., Walmsley, A., Vahldick, H.E., Rybka, D., Leite, M.F., Musa, Z., Bucksch, A., et al., 2024. The soil microbiome modulates the sorghum root metabolome and cellular traits with a concomitant reduction of striga infection. Cell reports 43.

Kim, M.J., Moon, Y., Tou, J.C., Mou, B., Waterland, N.L., 2016. Nutritional value, bioactive compounds and health benefits of lettuce (lactuca sativa l.). Journal of Food Composition and Analysis 49, 19–34. doi:10.1016/j.jfca.2016.03.004.

Kugelman, J., Allman, J., Read, S., et al., 2022. A comparison of deep learning u-net architectures for posterior segment oct retinal layer segmentation. Scientific Reports 12, 14888. doi:10.1038/s41598-022-18646-2.

Leite, D., et al., 2024. Advancements and outlooks in utilizing convolutional neural networks for plant disease severity assessment: A comprehensive review. Smart Agricultural Technology 9, 100573. doi:10.1016/j.atech.2024.100573.

Li, Z., Stylianou, A., Pless, R., 2019. Learning to correct for bad camera settings in large scale plant monitoring, in: IEEE Workshop on Computer Vision Problems in Plant Phenotyping (CVPR Workshop). URL: https://terraref.org/publication/li-2019-corrected-rgb-exposure.html.

Liu, S., Barrow, C.S., Hanlon, M., Lynch, J.P., Bucksch, A., 2021. Dirt/3d: 3d root phenotyping for field-grown maize (zea mays). Plant physiology 187, 739–757.

Liu, S., Bonelli, W.P., Pietrzyk, P., Bucksch, A., 2023. Comparison of open-source three-dimensional reconstruction pipelines for maize-root phenotyping. The Plant Phenome Journal 6, e20068.

Moazzam, S., et al., 2021. A patch-image based classification approach for detection of weeds in sugar beet crop. IEEE Access 9, 121698–121715. doi:10.1109/ACCESS.2021.3109015.

OpenCV team,. Contours in opencv (python) – getting started with contours. URL: https://docs.opencv.org/3.4/d4/d73/tutorial_py_contours_begin.html. accessed: 2025-04-10.

OpenCV Team,. OpenCV: Open Source Computer Vision Library. URL: https://opencv.org/. accessed: 2025-04-10.

Rizaldi, A.A., Gautama, E., Kamelia, L., 2023. Performance measurement of resnet-34 in convolutional neural network method for classification of mask type usage, in: Proceedings of the 2023 6th International Conference of Computer and Informatics Engineering (IC2IE), pp. 115–120. doi:10.1109/IC2IE60547.2023.10331025.

Sankaran, S., Mishra, A., Ehsani, R., Davis, C., 2010. A review of advanced techniques for detecting plant diseases. Computers and electronics in agriculture 72, 1–13.

Scott, J., Gordon, T., Kirkpatrick, S., Koike, S., Matheron, M., Ochoa, O., Truco, M., Michelmore, R., 2012. Crop rotation and genetic resistance reduce risk of damage from fusarium wilt in lettuce. California Agriculture 66, 20–24. URL: https://escholarship.org/uc/item/7s63x3t6.

Sudars, K., Namatevs, I., Nikulins, A., Balass, R., Peter, A., Strautina, S., Kaufmane, E., Kalnina, I., 2023. Semantic segmentation using u-net deep learning network for quince phenotyping on rgb and hyperspectral images, in: 2023 27th International Conference Electronics, pp. 1–4. doi:10.1109/IEEECONF58372.2023.10177638.

US Department of Agriculture, 2020. Usda: Agricultural marketing service market news. https://www.ers.usda.gov/data-products/chart-gallery/gallery/chart-detail/?chartId=106516.

V, S., Bhagwat, A., Laxmi, V., 2024. Leafspotnet: A deep learning framework for detecting leaf spot disease in jasmine plants. Artificial Intelligence in Agriculture 12, 1–18. doi:10.1016/j.aiia.2024.02.002.

Wang, X., Polder, G., Focker, M., Liu, C., 2024. Sága, a deep learning spectral analysis tool for fungal detection in grains—a case study to detect fusarium in winter wheat. Toxins 16, 354. doi:10.3390/toxins16080354.

